# SciReader: A Cloud-based Recommender System for Biomedical Literature

**DOI:** 10.1101/333922

**Authors:** Priya Desai, Natalie Telis, Ben Lehmann, Keith Bettinger, Jonathan K. Pritchard, Somalee Datta

## Abstract

With the growing number of biomedical papers published each year, keeping up with relevant literature has become increasingly important, and yet more challenging. SciReader (www.scireader.com) is a cloud-based personalized recommender system that specifically aims to assist biomedical researchers and clinicians identify publications of interest to them. SciReader uses topic modeling and other machine learning algorithms to provide users with recommendations that are recent, relevant, and of high quality^1^.

## I. INTRODUCTION

Today’s researchers have access to large online scientific repositories of literature and face the monumental challenge of sifting through the vast amounts of available information to find articles relevant to their interests. These digital archives are growing rapidly as new articles are put online and old publications are scanned, indexed, and made digitally available. While keeping up with the ongoing work in their discipline is an essential task for scientists, it can be tedious and time-consuming. Furthermore, with the growing number of publications, it is nearly impossible to access and track the most relevant and promising research articles. This points to the need for new technology solutions that enable researchers to access publications central to their disciplines, understand the current trends, and collect references for their own research. While, historically, researchers have found articles by following citations in other articles of interest, today’s fast-paced research world makes that method cumbersome and insufficiently comprehensive. Today, the most common strategies for finding relevant articles are through keyword-based web searches, Google Scholar searches, or by scanning through RSS feeds from popular journals. Sites like *CiteU-Like*[11] and *Mendeley*[17] allow researchers to create their own reference libraries for articles they are interested in and enable sharing with peers[13]. These shareable libraries and vast amounts of literature makes this field ripe for using a filtration system—and a good recommender system[9] (which filters out the least relevant articles) could be invaluable.

Recommender systems have long been widely used in e-commerce as valuable tools for personalizing purchase recommendations. *Amazon* and *Netflix* suggest products and movies based on each user’s profile, previous purchase history, and online behavior. *Facebook* and *Google* news recommend news articles based on user profiles and behavior. As the amount of information available grows, personalized recommendation systems have become crucial tools for finding useful content.

## II. BACKGROUND AND RELATED WORK

While an explosion of available scientific literature is occurring in many fields, we focus specifically on the area of biomedical literature (Fig 1). In the burgeoning field of biomedicine, a staggering 3,000-5,000 papers are published every day (Fig 2). It is not practical to browse through so many articles to identify those that may be relevant. For newcomers to the field, it is hard to establish a baseline understanding of seminal work. New journals are popping up on a regular basis[27]. Good recommender systems can play a valuable role in winnowing down publications of interest.There are primarily two kinds of recommender systems:

**Fig. 1.**
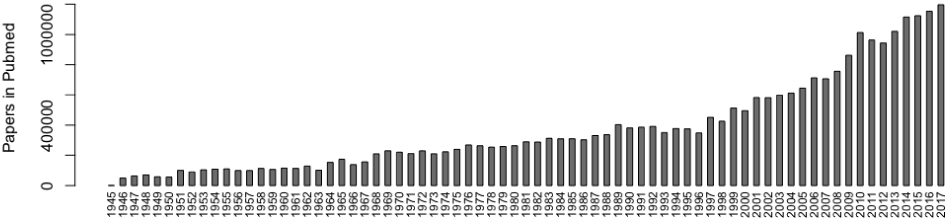
Number of Papers Uploaded to PubMed per Year.

**Fig. 2.**
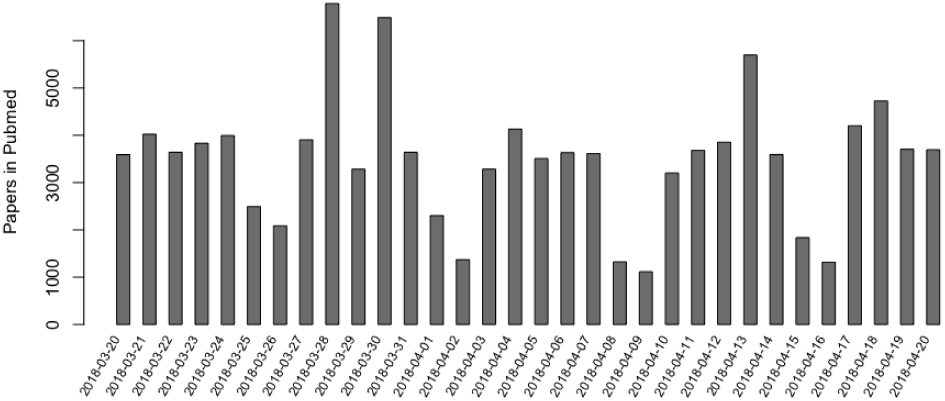
Number of Papers Uploaded to PubMed per day between 3-20-2018 and 4-20-2018.

1. Content-based recommender systems, and
2. Collaborative filtering-based recommender systems.

Content-based filtering methods are usually based on a description of the item and a profile of the user’s preferences. They utilize discrete characteristics (attributes) of an item in order to recommend additional items with similar properties.These algorithms recommend items that are similar to those that the user liked in the past. An important drawback of content-based filtering is that the system is unable to interpret user preferences in one context and use them across other content types. It tends to recommend only similar items and the recommendations have very little novelty factor. *Rotten Tomatoes*[14] and *Pandora Radio*[16] are popular content-based recommendation systems.

Collaborative filtering-based approaches[10] typically build models based on user profiles, user’s past behaviors (items previously purchased or selected and/or numerical ratings given to those items), and the behavior of similar users to recommend items[15]. A key advantage of the collaborative filtering approach is that it does not require a large number of item attributes. However, it suffers from the cold start problem – it typically requires a large amount of user data to make meaningful recommendations. *Last.fm*[30] is a music station that primarily uses the collaborative filtering approach.

Modern recommenders often follow a *hybrid*[6] approach, employing both content-based and collaborative-filtering-based techniques. *Netflix*[18] is a good example of a hybrid recommender system.

In recent years, a number of websites that can provide scientific recommendations have been developed: e.g. Scienstein[6], Google Scholar[32], PubChase[20], Mendeley[13], [17], Sparrho[31] and others. Many of these sites focus mainly on citation counts and classic text mining strategies, each of which has advantages and potential pitfalls, thus limiting its general usability. For example, citation databases are often incomplete, and have issues disambiguating homographs. They may also be subject to the Matthew Effect[21],which is the sociological phenomenon that eminent scientists will often get more credit than a comparatively unknown researcher, even if their work is similar. Text-based recommenders often have trouble with synonyms and context-based words, and often cannot identify papers that may be related.

## III. SCIREADER

### A. An Overview

SciReader is a hybrid recommender system and takes advantage of both the representation of the content as well as the similarities among users. While most biomedical recommender systems like PubChase focus on finding new items, we believe that sharing, discussing, and reviewing papers are an integral part of a scientist’s professional activities, and these motivations are reflected in our requirements for the SciReader site. Our general goals for SciReader are:

i. The articles recommended should be relevant to the user.
ii. The articles should be fairly recent. While we believe it is important to be able to recommend older relevant articles, our immediate motivation is to address the problem of current data deluge.
iii. Users should be able to view, comment, and share via email and social media, articles they find relevant and interesting.
iv. Users should be able to bookmark articles they like or plan to read and should be able to organize articles from different projects.
v. Twitter, which is increasingly used to announce new results and publications, should be integrated into SciReader.
vi. It should be to useful to newcomers in the field – not just established scientists with many publications. It should allow users to easily navigate the extensive body of biomedical literature.

The two main parts of this system:

i. Recommendation Algorithm
ii. User-system Interaction

are discussed in the following sections. Fig 3 below is an overview of the SciReader recommender system.

**Fig. 3.**
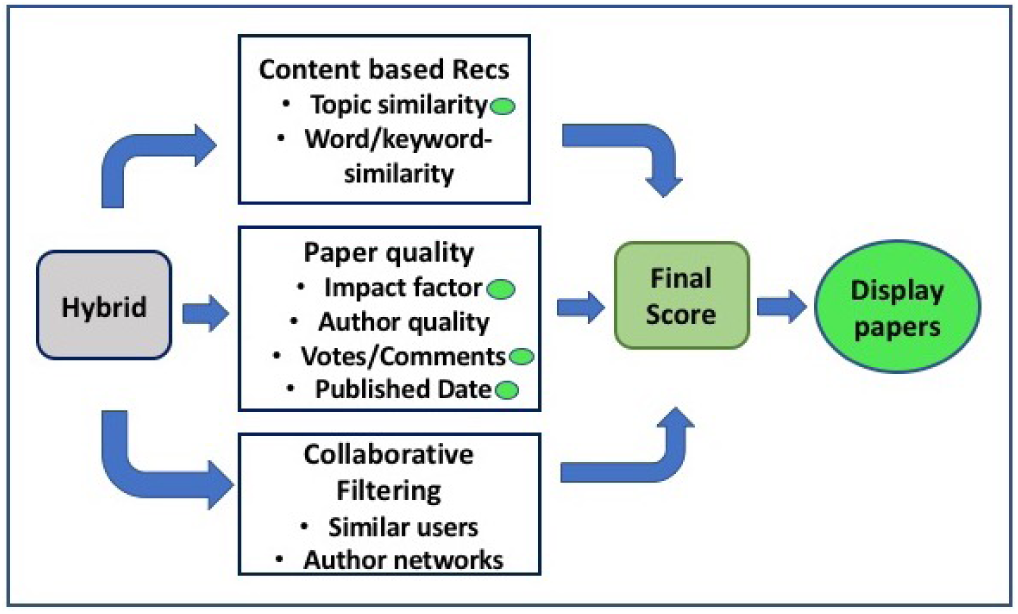
The Recommendation system:An Overview

### B. Recommendation Algorithm

Generating recommendations is a multi-step matching and filtering process that combines multiple algorithmic approaches as described below.

1. *Probabilistic Topic Models:* Topics can be thought of as “recurring patterns of co-occurring words”[28], and a topic model is a type of statistical model often used to discover abstract topics that occur in a collection of documents[1], [2]. Topic models are frequently used to discover hidden semantic structures in a large text body[7], [8]. One of the most commonly used topic models is latent Dirichlet allocation (LDA)[2]. LDA is a matrix factorization technique and assumes documents are produced from a mixture of topics. Unlike a clustering algorithm, where each document would be assigned to one cluster, LDA allows documents to belong to multiple topics with varying probability. Given a dataset of documents, LDA backtracks and tries to figure out what topics would create those documents in the first place. LDA specifies a *generative process*, an imaginary probabilistic recipe that produces both the hidden topic structure and the observed words of the texts. Those topics then generate words based on their probability distribution. For example: Assume there are *K* topics *β*= *β*_1:*K*_, each of which is a distribution over a fixed vocabulary. The generative process of LDA is as follows: For each article *w*_*j*_ in the corpus,
  1. Draw topic proportions *θ*_*j*_ ∼ Dirichlet(*α*).
  2. For each word *n*, (a)
    a. Draw topic assignment *z*_*jn*_∼ Mult(*θ*_*j*_).
    b. Draw word 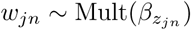.
  This process reveals how the topic proportions are document-specific, but the set of topics is shared by the corpus. Topic modeling algorithms perform what is called probabilistic inference. Given a collection of texts, they reverse the imaginary generative process to answer the question: “What is the likely hidden topical structure that generated my observed documents?” The posterior distribution (or maximum likelihood estimate) of the topics reveals the K topics that likely generated its documents [23], [24].
2. *PubMed Topic Model:* The cornerstone of SciReader is its topic model of PubMed. PubMed (http://www.ncbi.nlm.nih.gov/pubmed) is a free search engine maintained by the NIH primarily accessing the MEDLINE database of references and abstracts on life sciences and biomedical topics. It provides a fairly comprehensive coverage of all articles published in biomedicine. Metadata information about each article, including authors, title, institution, and abstract, is freely available. We used titles, abstracts, and keywords from all articles available in PubMed and published in 2012 to create our document collection (or text corpus) and then generate topics. Only abstracts (not full text) were chosen because they are freely available for all articles. Full texts are not available for most publications. We use MALLET (Machine Learning for Language Toolkit)[5], [22], a Java-based package for statistical natural language processing developed at the University of Massachusetts at Amherst, to run an implementation of LDA. The corpus from PubMed that we used contained approximately one million unique articles, and for each article, we concatenated its title, abstract, and keywords. We applied heuristics to clean up the data, like removing standard English stop words (like “a”, “do”, and “these”)and word stemming. To generate topics, we used the vector space model representation of the corpus[33]. The final training corpus had approximately 650,000 distinct words and was used to generate a topic inferencer which classified biomedical papers into 150 topic areas. The number of topics was determined after much manual parameter adjustments and based on multiple factors including Silhouette clustering[26] and perplexity[25]. For ease of use, we manually examined the LDA output to provide an informative name for each automatically defined topic. This inspection was done by examining the most frequent words in the topic, the typical journals where articles with a high probability of this topic were published, and the actual articles containing a high probability of that topic. Some topics which seemed to capture a large number of non-english words and had a noticeably fewer documents with high probabilities were dropped in the final topic display(Fig 7a). Fig 4 displays the most frequent words of a topic as a word cloud along with the curated topic name. The size of the word in the word cloud is proportional to the relative frequency of the word in that topic. The topic names are human-curated.

**Fig. 4.**
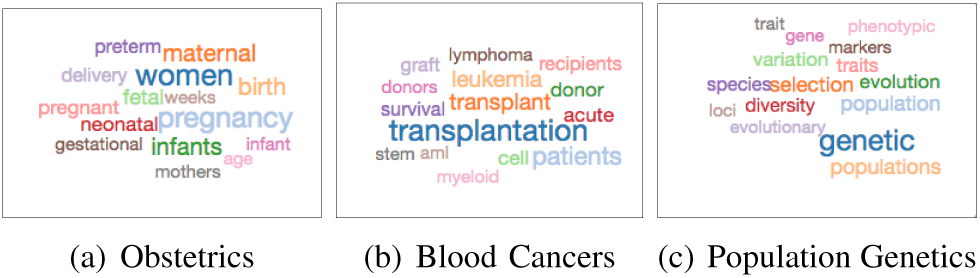
Examples of Topics generated by LDA: For better navigation on the website, we further grouped these topics manually into twenty *supertopics*. Fig 5 is an example where *Genetics and Genomics* is the supertopic created to contain seven sub-topics, each of which were generated by the algorithm.

**Fig. 5.**
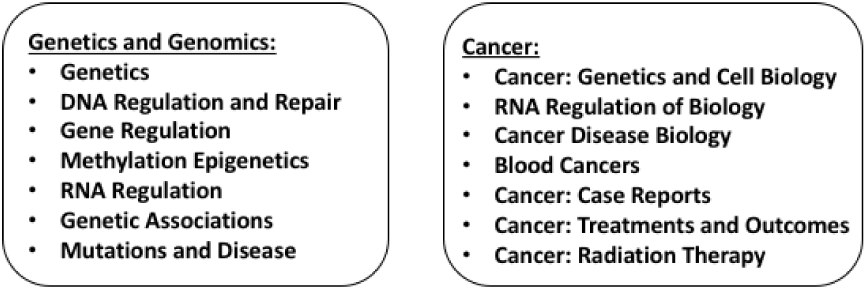
Supertopics and Corresponding sub-topics The topic inferencer was stored and is used to infer topics for all new documents. All the older articles in PubMed have also been processed using this inferencer, and stored in terms of their topics. Thus, each paper is classified into one or more topics by using the same LDA inferencer model with fixed topic definitions. This topic model can be considered to be an interpretable, low-dimensional representation of PubMed and we exploit this representation to find *topically similar* papers.
3. *Tf-idf and cosine similarity:* As part of preprocessing, all articles or documents are converted into a vector space representation matrix. The documents are tokenized using the same *≈* 650,000 word dictionary that was generated while creating the LDA inferencer. We then apply the tf-idf normalization[34], [35] to the matrix of documents and use that representation to calculate the cosine distance between the documents. The tf-idf weight is a statistical measure used to evaluate how important a word is to a document in a collection or corpus by assigning a higher weight to rarer words in a collection of documents. It can be expressed as:

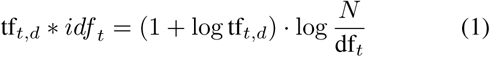

where *tf*_*t,d*_ is the number of occurrences of term *t* in document *d*, *N* is the total number of documents in a collection and *df*_*t*_ is the document frequency of term *t*. Document similarity or distance between documents is a standard method used in information retrieval to measure how semantically similar two documents are[33], [35]. While there are different distance metrics (Jaccardean, Euclidean, Manhattan) that can be used, cosine measure, which is the Euclidean dot product between the vector representation of the two documents, is frequently used as a measure of document similarity and the metric that gave us the best results.

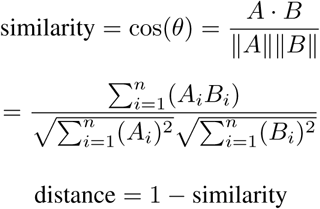 The *smaller* the distance, the more *similar* the documents. To make recommendations, we generate matrices: one using the articles in the user’s library, and the other using the articles in the corpus, and then we find the papers most similar to the user’s library in the corpus.

### C. User-System Interaction

Since SciReader is focused on providing personalized recommendations in the biomedical literature space, we need to gather a variety of different kinds of information about the user to understand their specific research interests. When a user initially registers with the site, they are led through a series of questions and asked to provide at least one of the following:

1. topics of interest from the predefined list of *≈* 150 topics. These are the topics generated by the topic modeling algorithm as discussed above (Fig 7b).
2. keywords that describe their research interests(entered as free-format text) (Fig 7b).
3. authors of interest.
4. an existing reference library/bibliography (Fig 7c).
5. a personal publication list (Fig 7c).

All users are provided with a couple of basic libraries: *Reading List* and *My Publications*. These libraries are meant to serve as placeholders for the users (yet to be populated) reading list and publications. As useful papers from the recommendation list are found, they can be added to the reading list. The users are free to create as many more libraries as they need to organize their recommendations and references. The site has a built-in interface with PubMed, so a user can easily search for their own and other publications and add them to their libraries. The central idea is that newcomers to the field can get a general flavor of the current publications by providing only topics and keywords of interest. They can then peruse the recommendations by topic which are updated daily, and slowly build their digital library. Established practitioners of the field who may already have personal reference libraries can upload them right away (Fig 7c) in addition to providing basic keywords and topics. All this information gives the recommender more data to work with and helps provide more personalized recommendations.

A user can interact with the site by liking, upvoting, sharing or commenting on an article. These modes of feedback provide the recommender with valuable information about the user’s current interests. Our philosophy is that a small amount of initial information can assist us with identifying general areas of interest, but after that, the user’s behavior within the site will be most helpful for fine-tuning the recommendations. For example, we encourage users to create separate libraries for different projects, because separate libraries allow the recommender to tailor its recommendations to each project. Recommendations are created for each individual library and an *Overall* recommendation set is compiled of the recommendations from all the libraries. SciReader tracks every time a recommended paper is uploaded into one of the user’s libraries and records all interactions the user has with the site.

In addition to paper recommendations, SciReader also provides users with regular Twitter updates relating to papers in their topics of interest. We monitor Twitter accounts from most prominent journals and publications and crosslink them with articles from PubMed. Tweets that refer to publications are then displayed as part of the users recommendation feed. Fig 6 describes the typical user workflow and Figs 7(a-c) show different aspects of the user portal.

**Fig. 6.**
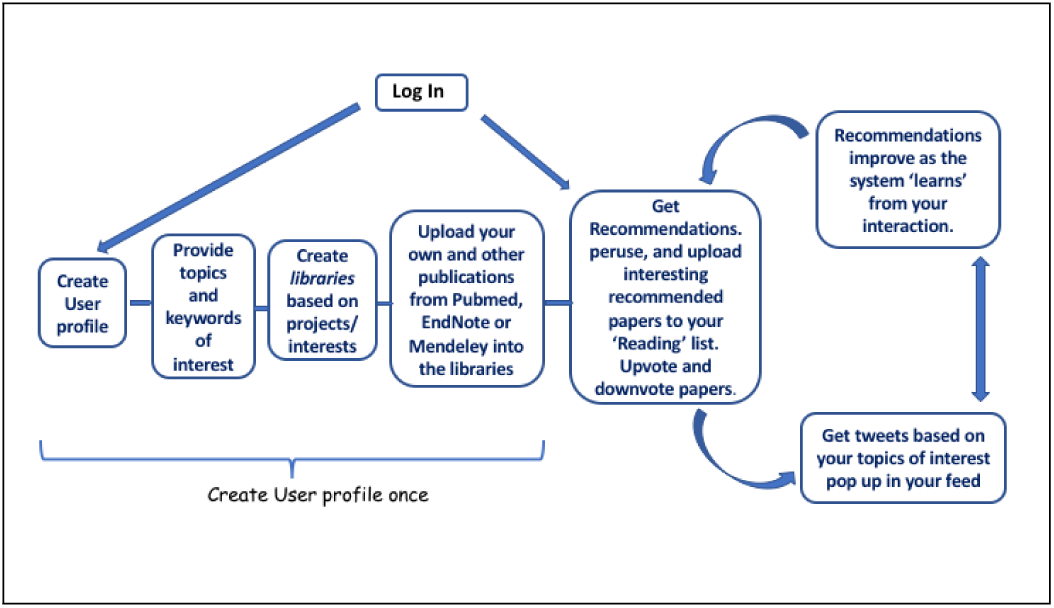
Typical user workflow.

**Fig. 7.**
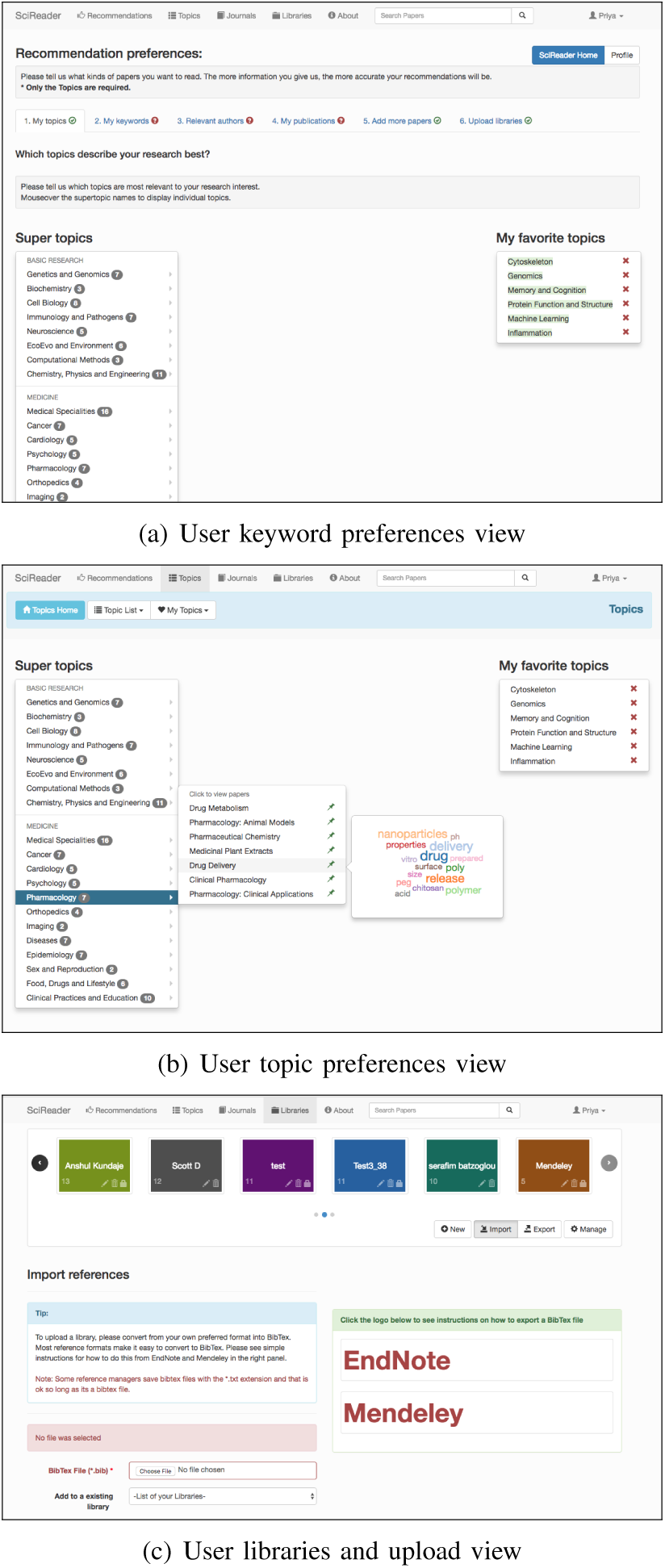
Different facets of the User portal

### D. Finding relevant papers

We assume that a user *k* has a library *L* of *n* papers, and that we want to compare these against a corpus *C*,containing *m* new papers. We use a filtering step to prepare recommendation set *R*, in which we only include papers in *C* if they have substantial topic overlap with papers in *L*. This intersection is done by comparing the topic representations of papers in the user’s library versus the corpus. Next, we compute a word similarity matrix *W*_*ij*_, between each paper *∈ l*_*I,*_ *L* and *c*_*j*_, *∈C*. The entries in *W* are in [0, 1] with 1 indicating complete similarity, and 0 indicating complete difference. We then identify papers in *C* that are especially close to at least some papers in *L*. We use the following distance function to assign a score to each paper *c*_*j*_, as follows:

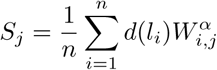

where *d*(*l*_*i*_) is a function of the date of publication of paper *l*_*I*_ such that similarity to older papers in the library is down weighted (since those may reflect less recent user interests), and *α≤* 1 is a constant that upweights high similarities. Our rationale here is that medium values of *α≤* 1 result in upweighting of papers which are overall similar to clusters of papers in the library, rather than picking either papers that have similarity to everything but not very similar to anything, or papers that are extremely similar to one paper only.

The impact factor of a journal is frequently used as a proxy for the relative importance of a journal within its field. We upweight papers from journals with higher impact factors, as well as journals which published the papers in the user’s library and contain the keywords provided by the user. Furthermore, we would like our recommendations to be strongly biased towards more recent papers. We have developed a function that models these characteristics which we use to rank papers. ‘’

Mentions of papers in social media such as Twitter or blogs can be a leading indicator of the future importance of a paper. We search Twitter accounts for URLs that link to scientific papers and match these to papers in our database. For each paper, we record all tweets that are positively matched as referring to that paper. These data are currently summarized simply as the number of tweets referring to any given paper, but, in the future, we will consider more complicated weighting schemes, such as estimates of the tweeter quality based on past performance at identifying good papers.

We are further refining our prediction algorithm to incorporate a variety of features to predict how important a paper is likely to become. Currently, the key features used by our prediction algorithm include journal impact factor and Tweet+Like counts. Citation counts are frequently used as a measure of the importance of papers. However, it usually takes more than a year before a paper starts to accumulate citations, rendering this approach inapplicable for recommending new literature. We are working on developing an author and institution quality score based on citation count data and author position.

### E. Putting it all together: Creating recommendations

We routinely download new papers from PubMed to a local database. All new papers are then processed through the LDA inferencer and stored in a separate table. As part of pre-processing, a *current corpus* matrix of all the papers published the last 90 days is generated, using both the topic-model representation, as well as the tf-idf normalized form. There are typically between 275,000 - 400,000 papers in the current corpus. The dictionary used to tokenize the papers is the same one that was used in generating the LDA inferencer and is thus common for all papers. This *current corpus* is then used to generate all the daily and immediate recommendations.

All the user libraries are similarly processed and stored using the topic-model representation as well as the tf-idf normalized matrix representation using the same word vocabulary (dictionary). Whenever a user adds/removes papers from any of their libraries, or creates a new library, the matrices are updated accordingly. When a user uploads a new library, the *immediate recommendations* function is triggered. The user library is pre-processed and topic comparison is done between the library papers and current corpus. We filter out papers from the current corpus that don’t pass a minimum threshold of similarity with the user’s library, thus ensuring that only topically similar papers are further considered for recommendation. We then compute the word-similarity matrix between the user’s library and this *filtered current corpus* and generate a score for each paper. The top *n* (typically a few hundred) papers are taken and only these will be used to generate the final recommendations. These *n* papers are then re-scored using the weighting function that factors in the date of publication, journal impact factor, journals included in the user’s library, keywords, etc., and re-ranked. The papers are sorted by this new ranking and then displayed as the user’s recommendations. Overall recommendations are a compilation of recommendations from all the libraries for a specific user, re-ranked using certain criteria. The process for generating daily recommendations is similar except that we run this recommendation for all libraries and for all users. Fig 8 shows a schematic workflow and Fig 9 shows a screenshot of a typical users library recommendations.

**Fig. 8.**
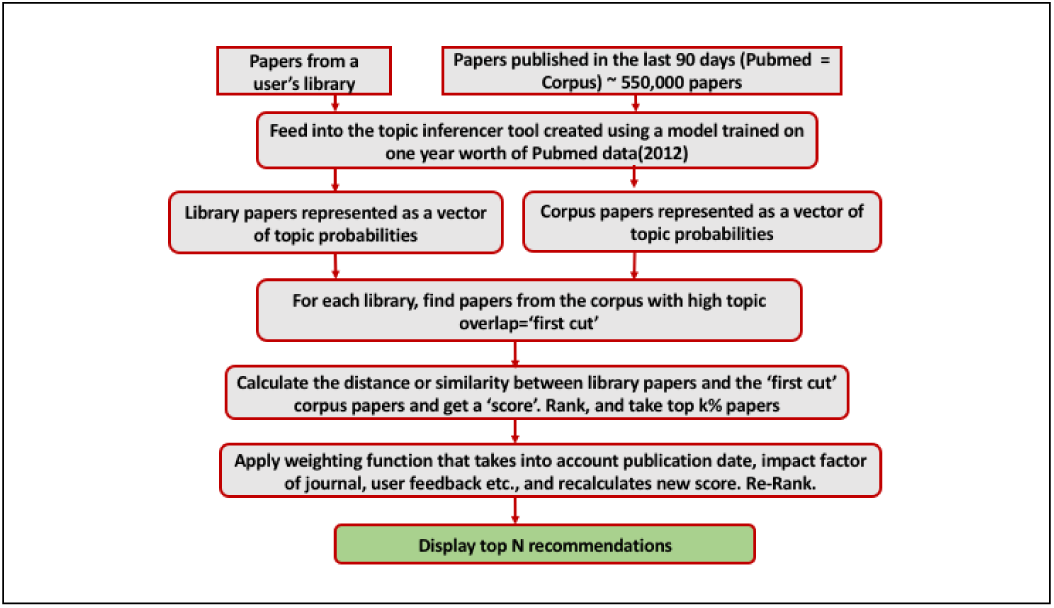
Recommendation workflow

**Fig. 9.**
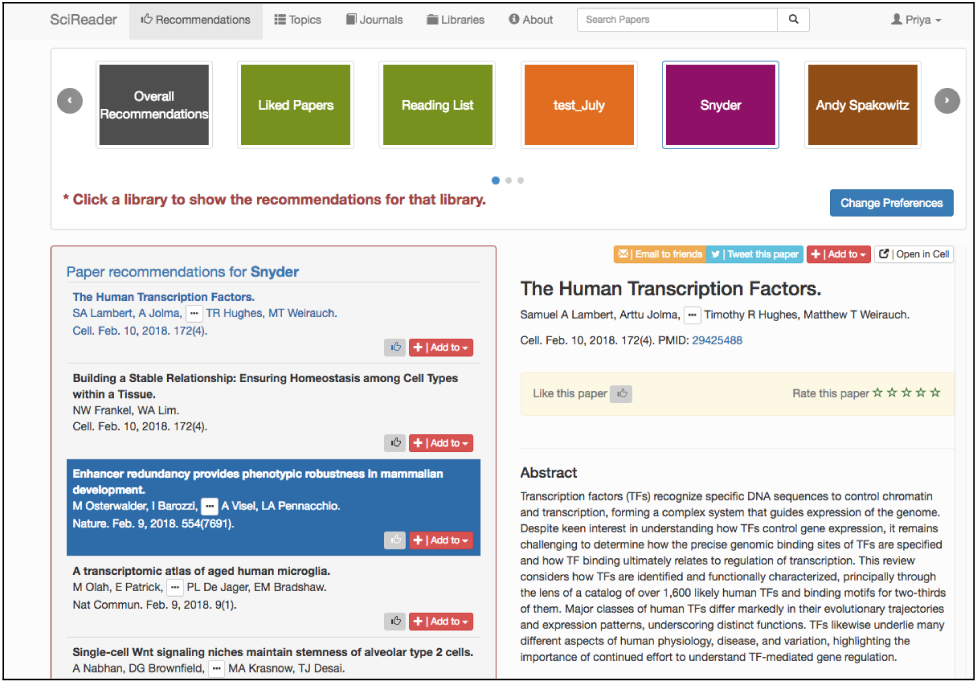
Individual library recommendations.

In addition to paper recommendations, we also provide a daily updated tweet feed. As stated earlier, we have created a database linking tweets and the digital object identifier (*doi*) of the publications and keep track of how many times a publication is tweeted or re-tweeted about. Since the publication has been through our topic inferencer, we know the main topics in the publication, and thus can assign the tweet the corresponding topic(s). The tweet is thus labeled as belonging to a certain topic. The tweet feed is a list of tweets that refer to papers that refer to the users preferred topics and had the highest number of tweets. Tweets that do not refer to a specific topic, but are tweeted about very frequently are labeled as “General”. Thus, each user gets a Twitter feed uniquely personalized to them. Fig 10 shows a typical Twitter feed.

**Fig. 10.**
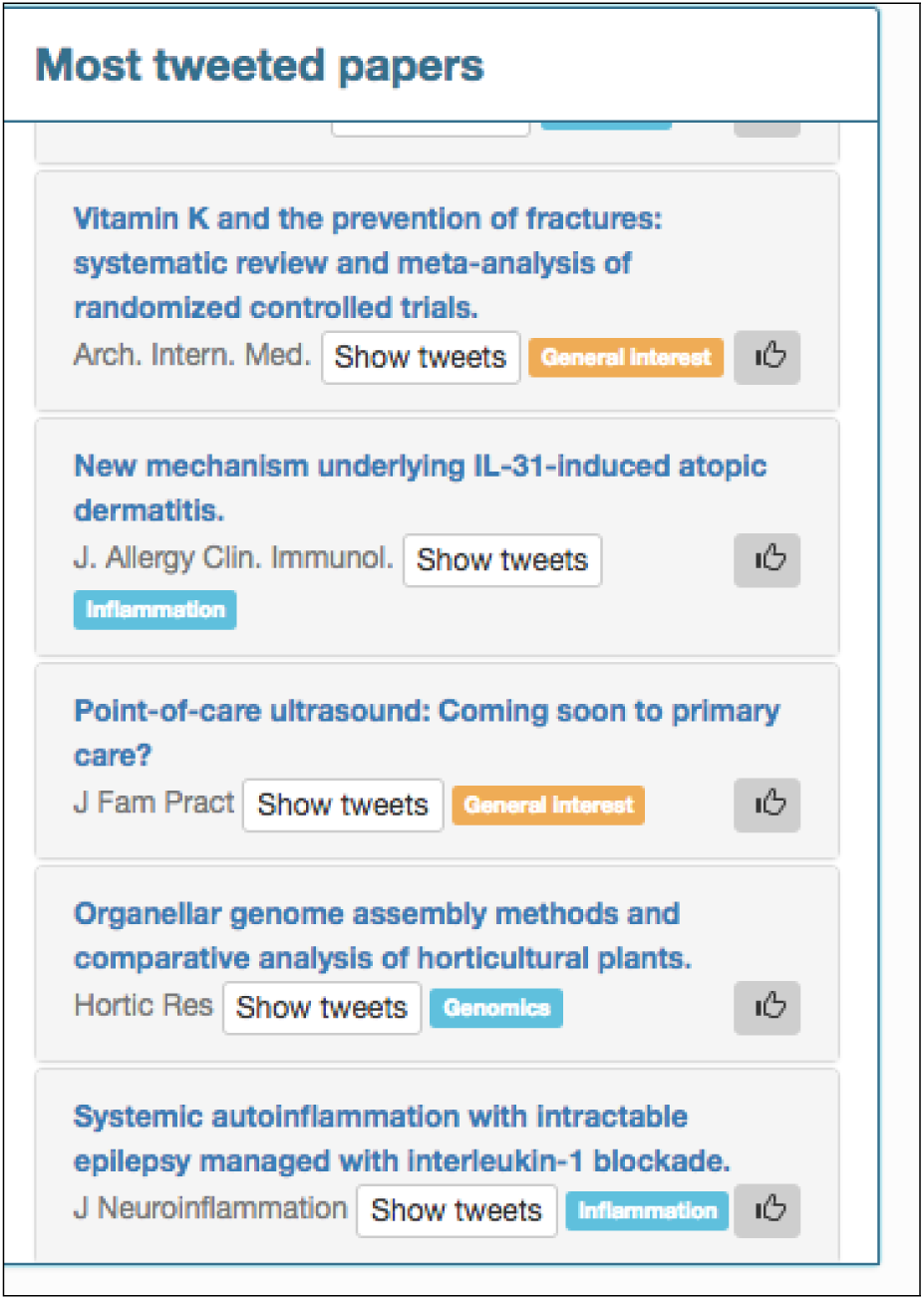
Screenshot of a user’s Twitter feed.

We also generate a weekly digest of recommendations which emailed to each user. This digest is comprised of the papers that were at the top of their recommendation feed for that week. All the data processing on the backend is done using python and the web interface uses the Django 1.7 framework.

### F. Optimizing Calculations

Providing on demand, instantaneous recommendations is computationally intensive and input/output of data can time-consuming. The actual word-similarity calculation often involves multiplications between matrices of size *≈* 650,000 × 15,000 and, reading these in and performing the calculation can be both computationally and memory intensive. However, since our vector space model matrices are sparse, we can use the sparse matrix format to significantly reduce the memory footprint. Hierarchical Data Format version 5 (HDF5) [36] is a great mechanism for storing large quantities of numerical data which can be read in rapidly and allow for sub-setting and partial I/O of datasets. We use the H5py package/PyTables, a pythonic interface to the HDF5 binary data format which allows easy manipulation of data. The sparse tf-idf matrices are stored in the hdf5 format and calculations done using Scipy are coded in C/C++, thus significantly speeding performance. cPickle written in C is a python module that implements a fundamental, but powerful algorithm for serializing and de-serializing a Python object structure and is used to store the word token dictionary. We pre-calculate as many of the computations as possible so the recommendations can be generated efficiently.

**Fig. 11.**
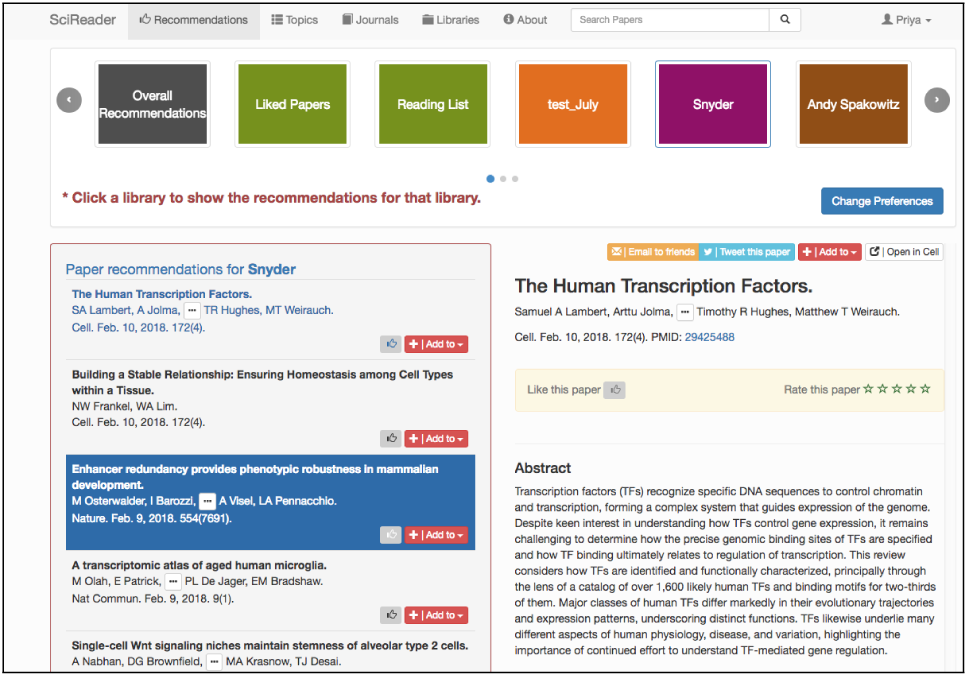
Library recommendations for a specific user.

### G. Cloud Architecture

SciReader was initially developed on Amazon Web Services (AWS) but has since been migrated to the Google Cloud Platform. Fig 12 shows the system architecture on Google Cloud. The user data as well as the PubMed metadata is stored in a MySQL database on the cloud. Google buckets store the precomputed matrices and the computations are done on a Google Compute Engine virtual machine. The website is hosted on a small virtual instance and an elastic load balancer is used to spin up more instances based on demand.

**Fig. 12.**
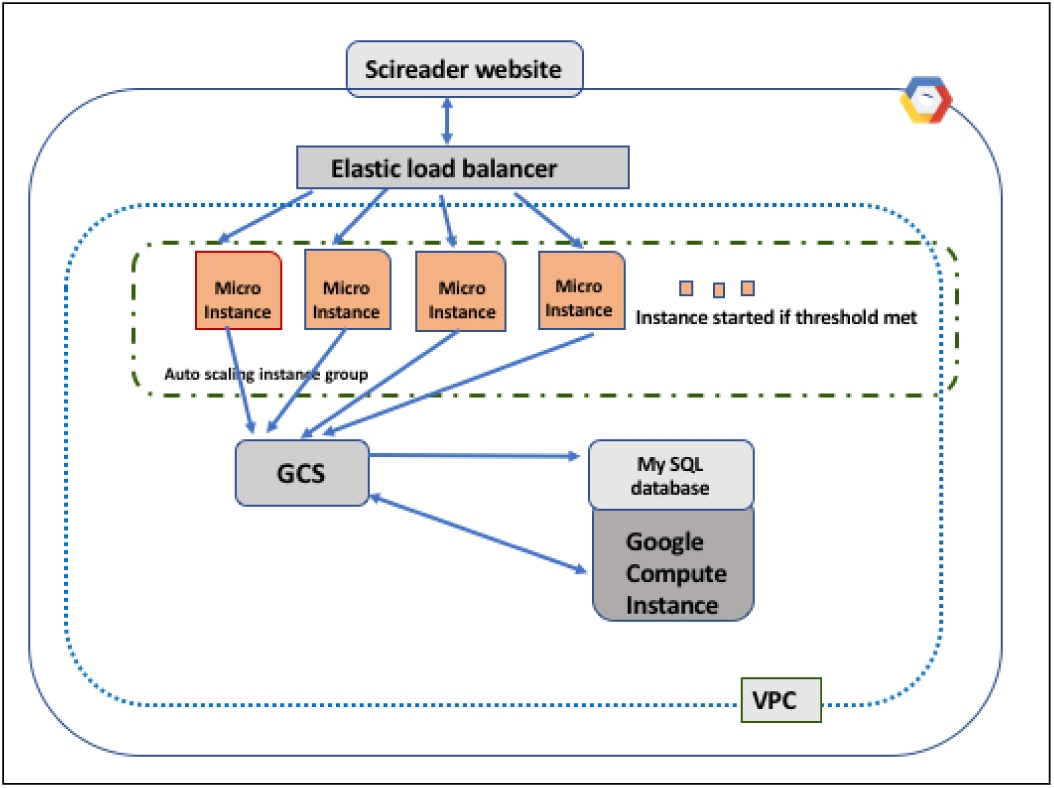
A schematic diagram of the current SciReader architecture on Google Cloud.

### H. Discussion and Future Work

SciReader is a fully functional recommender system for biomedical literature and it can be accessed at http://www.scireader.com. It currently has *≈* 1500 unique active users. We have conducted multiple informal focus groups to better understand our user base, and based on the feedback, we are working on improving and adding certain features and functionalities to the site. We are also planning on adding content from NIH RePORTER, arXiv, and bioRxiv to the recommendations. Further, we are developing a SciReader API so that the highly annotated database containing the topic representation for all biomedical articles used by SciReader can be accessible to other researchers for future bibliometric and longitudinal studies. In addition to improving the speed to generating recommendations, we hope to provide *user-tunable recommendations* where the user could explicitly choose which criteria to use to generate their personalized recommendations.

## ACKNOWLEDGMENTS

This project was initially developed in the Pritchard Lab and supported by funding from Stanford University and the Howard Hughes Medical Institute. It is now maintained by SCGPM. We thank Yonggan Wu for his help in building the SciReader website.

## Footnotes

* www.scireader.com

1 SciReader codebase will soon be released as open-source code.

